# High prevalence of Ethiopian and Indo-Oceanic *Mycobacterium tuberculosis* lineages among pulmonary tuberculosis cases in East Gojjam, Northwest Ethiopia: insights from genomic characterization

**DOI:** 10.64898/2025.12.21.695839

**Authors:** Kelemework Adane, Zelalem Desalegn, Elena Hailu

## Abstract

**Introduction:** *Mycobacterium tuberculosis* (Mtb) strains in Ethiopia exhibit significant genetic and regional variation, highlighting the need for further research across diverse geographic areas.

**Objective:** We employed a molecular tool to characterize Mtb isolates from predominantly rural patients at two public hospitals in East Gojjam, Amhara Region, northwest Ethiopia.

**Methods:** Molecular typing, including the region of difference 9 (RD9)-based polymerase chain reaction (PCR) and spoligotyping, was performed on stored sputum samples at Armauer Hansen Research Institute (AHRI) in Addis Ababa.

**Results:** Out of 120 Mtb, 116 yielded interpretable spoligotyping results, revealing four major lineages. The Euro-American (Lineage 4) was most common (42.6%), followed by the Ethiopian (Lineage 7, 20.8%), Indo-Oceanic (Lineage 1, 20.0%), and East African-Indian (Lineage 3, 16.5%). The dominant sublineages were ETH1 (Lineage 7, 20.8%), T (Lineage 4, 15.6%), and CAS1-Delhi (Lineage 3, 13.9%). In total, 39 spoligotype patterns were identified, including 19 shared types (48.7%) comprising 90 strains from the SITVIT2 database. The most frequent spoligotypes were SIT 910 (23.3%) and SIT 53 (14.4%), followed by SIT 54 and SIT 149 (10.0% each). Overall, 82.7% of the strains were clustered.

**Conclusion:** This study reveals substantial genetic diversity among Mtb strains in a rural area of Ethiopia, with a notably high prevalence of Lineage 7 and Indo-Oceanic strains. The high clustering rate (82.7%) suggests ongoing local transmission, particularly involving the ETH1, CAS1-Delhi, and Manu2 sublineages. Due to the distinct strain distribution, whole-genome sequencing is essential to better understand Mtb diversity and inform targeted TB control strategies.

## Introduction

Despite being both curable and preventable, tuberculosis (TB) remains the leading cause of death from a single infectious agent worldwide. In 2023, an estimated 10.8 million people developed active TB, and 1.25 million died from the disease (1). TB is caused by a genetically related group of bacterial species collectively known as the *Mycobacterium tuberculosis* complex (MTBC). These include more than eight species, such as *Mycobacterium tuberculosis* (Mtb) and *Mycobacterium bovis*, but the vast majority of human TB cases are attributable to Mtb (2).

Over time, Mtb has evolved to withstand host immune defenses, adapt to varying environmental conditions, and respond to numerous selective pressures. As a result, it now exists in multiple genetically distinct genotypes. Identifying the circulating Mtb strains in specific regions is crucial for understanding TB epidemiology and deciphering transmission dynamics, both of which are essential for effective and timely TB control (3). Mtb strains and lineages can differ in key characteristics such as pathogenicity, transmissibility, propensity for developing drug resistance, and immune evasion strategies (4). Several molecular methods have been employed to define the population structure of Mtb, including spoligotyping of clustered regularly interspaced short palindromic repeats (CRISPR), analysis of mycobacterial interspersed repetitive units-variable number tandem repeats (MIRU-VNTRs), and whole genome sequencing (5, 6).

To date, nine distinct lineages of Mtb have been identified globally, each showing unique geographic distributions and associations with specific human populations (7). These include: Lineage 1 (Indo-Oceanic, IO), Lineage 2 (East Asian/Beijing), Lineage 3 (East African-Indian, EAI), Lineage 4 (Euro-American, EA), Lineage 5 (*M. africanum* West African type 1), Lineage 6 (*M. africanum* West African type 2), Lineage 7 (Ethiopian), and the more recently discovered Lineages 8 and 9, found in East Africa and Uganda (7, 8). In general, the EA lineage is the most widely distributed and genetically varied, while the EA/Beijing lineage is highly virulent and frequently associated with drug resistance, with significant global coverage. Other lineages are relatively regionally confined and affect specific populations.

Ethiopia ranks among the 30 countries with the highest burdens of TB and TB/HIV co-infection, with an estimated TB incidence rate of 146 cases per 100,000 population and 19 TB-related deaths per 100,000 in 2023 (1). Although Ethiopia has recently been removed from the list of countries with a high burden of drug-resistant TB, about 54% of drug-resistant TB cases remain undetected each year. Compounding this challenge, recent national crises—including armed conflict, drought, and the impacts of COVID-19—have significantly hindered progress in TB control (2). Given the country’s large population and high TB transmission rates, molecular epidemiological studies are critical for revealing transmission patterns and guiding control strategies. A recent systematic review identified five main Mtb lineages circulating in Ethiopia: EA (64.8%), EAI (23.0%), IO (7.1%), Ethiopian (4.1%), and East Asian /Beijing (0.2% (9). However, genetic diversity and the geographic distribution of these lineages vary widely across the country, indicating a need for more localized research.

In this study, we characterized Mtb isolates obtained from predominantly rural TB patients attending two public hospitals in the East Gojjam zone of the Amhara region in northwest Ethiopia. We used spoligotyping, a PCR-based method that detects genetic polymorphisms based on the presence or absence of 43 known spacer sequences within the direct repeat (DR) locus of the Mtb genome. Although spoligotyping has relatively low discriminatory power compared to more advanced methods such as MIRU-VNTR and whole genome sequencing (10–12), it is a simple, rapid technique with access to large international databases. For these reasons, it remains one of the most widely used genotyping methods for Mtb (13).

## Materials and Methods

### Study setting and source of the isolates

The Mtb strains were recovered from presumptive TB patients attending two public hospitals in the East Gojjam Zone of northwest Ethiopia. The zone is divided into 16 rural districts and 4 urban administrations, with the administrative center at Debre Markos. It has 10 public hospitals, including one referral hospital and nine primary hospitals, 101 health centers, and 430 health posts. We collected sputum samples from 385 adult presumptive TB cases attending Motta General Hospital and Debre Markos Referral Hospital between 2011 and January 2012 **(Fig 1).**

**Fig 1.**
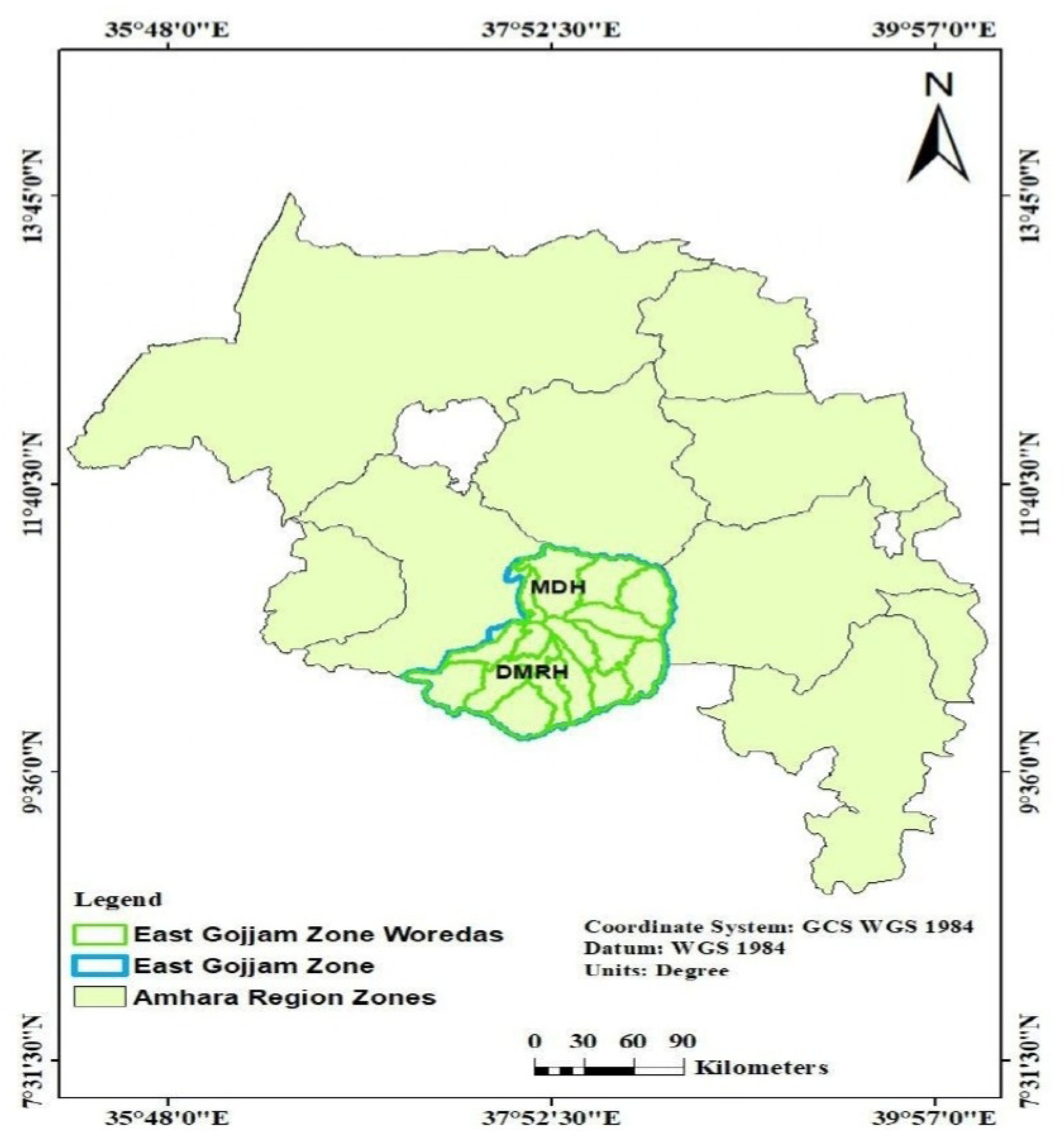
Map showing the study zone and hospital locations. MDH: Motta District Hospital; DMRH: Debre Markos Referral Hospital. Source: Generated by the researcher using the Amhara region shapefile (2025).

We also collected important patient characteristics, including sociodemographic features and clinical characteristics, using a structured questionnaire. A total of 124 Mtb isolates were collected. While the drug susceptibility profiles of these strains were published in 2015 as mentioned above (14), genotyping had not been completed until recently. The current study was therefore conducted on stored (archived) samples at the AHRI laboratory. We accessed the archived mycobacterial isolates collected on November 15, 2022, after which laboratory investigations were conducted.

### Mycobacterial culture and species identification

The procedures for mycobacterial culture and identification were detailed in our previous publication (14). Briefly, samples were decontaminated using Petroff’s method with 4% NaOH and cultured on Lowenstein-Jensen (LJ) solid media by standard protocols. Mycobacterial isolates were further characterized using various methods. Species identification was performed through RD9 deletion typing on heat-killed suspensions, utilizing FlankFW, RD9 Int, and RDFlankRev primers, as previously described by Brosch et al (15). A 396 bp PCR product indicated the presence of RD9, which is specific to Mtb and absent in other members of the Mtb complex.

### Molecular typing using Spoligotyping

A total of 120 MTBC isolates were spoligotyped using the method described by Kamerbeek et al (12). The direct repeat (DR) region was amplified using primers derived from the DR sequence: DRa (5′-GGTTTTGGGTCTGACGAC-3′) and DRb (5′-CCGAGAGGGGACGGAAAC-3′). PCR products were denatured at 96°C for 10 minutes and hybridized for 60 minutes at 60°C with a membrane containing 43 immobilized oligonucleotides, each corresponding to a unique spacer sequence in the DR locus. Following hybridization, membranes were washed and incubated with diluted streptavidin-peroxidase (HotStar, Crawley, UK) for 45–60 minutes at 42°C. DNA detection was carried out using enhanced chemiluminescence (ECL; Amersham Biosciences, UK), and the signal was visualized on X-ray film (Hyperfilm ECL, Amersham). Films were developed in a darkroom, fixed, dried, and analyzed.

The resulting patterns—black and white squares indicating the presence or absence of spacers—were converted into binary format (1/0) for subsequent analysis. DNA from reference strains *M. bovis* SB1176 and Mtb H37Rv served as positive controls, while water (Qiagen, Germany) was used as a negative control.

### Data analysis and Mtb strain retrieval

Data were recorded using a pre-designed Microsoft Excel spreadsheet. Statistical analysis was performed using SPSS software, version 28.0. Associations between variables were assessed using the chi-square test or Fisher’s exact test, as appropriate. Spoligotype patterns from the Excel spreadsheet were converted into binary and octal formats using the SITVITWEB tools, and Mtb strains were retrieved from various online sources. The SPOLDB4 and SITVITWEB databases (http://www.pasteur-guadeloupe.fr:8081/SITVIT2/batch.jsp) (16) were used to assign Shared International Types (SIT) numbers and identify related lineages. Patterns found in two or more global isolates were designated as SIT, while unique, unmatched patterns were classified as orphan strains. For further lineage classification, the RUN TB-lineage (https://tbinsight.cs.rpi.edu/run_tb_lineage.html) (17) and SpolLineages (http://www.pasteur-guadeloupe.fr:8081/SpolLineages/spol.jsp) tools were used to determine Conformal Bayesian Network (CBN) and SNP-based lineages, respectively. Strains sharing identical spoligotype\ patterns were defined as “clusters,” whereas distinct patterns observed in only one strain were considered “unique”. Furthermore, the MTBC lineage and sub-lineages were geographically mapped using ArcGIS version 10.8.

### Ethics Considerations

This study was conducted exclusively on archived mycobacterial isolates stored at the AHRI laboratory, with no direct patient contact and no new sample collection. Accordingly, the study was granted a waiver of ethical approval, as it involved the use of previously collected and anonymized laboratory samples only.

## Results

### Characteristics of the study population

The study population has been described in detail previously [14]. In summary, 124 MTBC isolates were obtained from sputum cultures of 385 individuals with suspected pulmonary TB. Forty-seven (37.9%) of these isolates were smear-negative, while 77 (62.1%) were smear-positive. Among the culture-positive participants, 82 (66.1%) were male and 42 (33.9%) were female, with a mean age of 31 ± 13 years. Multiplex PCR targeting RD9 confirmed 120 isolates as Mtb species. While the drug resistance profiles of these isolates have been reported earlier, this study focuses on their molecular diversity. Among the 89 isolates tested for drug susceptibility, 18 (20.23%) exhibited resistance to at least one anti-TB drug—12 (20.7%) from males and 6 (19.4%) from females (14).

### Genetic Diversity of Mtb Strains

#### Lineages and sublineages

Table 1 summarizes the spoligotyping results, including the lineages and sublineages of the 116 samples with interpretable spoligotyping results. We identified four major lineages among 115 Mtb isolates. The EA lineage (Lineage 4) was the most prevalent, comprising 49 isolates (42.6%). This was followed by the ETH (Lineage 7) and the ancient IO lineage (Lineage 1), with 24 (20.8%) and 23 (20.0%) isolates, respectively. The EAI lineage had the lowest representation, with 19 isolates (16.5%). The lineage of one isolate could not be determined. With regard to sublineages, the ETH sublineage of Lineage 7 was the most prevalent, accounting for 20.8% of the isolates. This was followed by the ill-defined T sublineage of the EA lineage, which included 18 out of 115 isolates (15.6%). The CAS1-Delhi sublineage of the EAI lineage ranked third, representing 16 isolates (13.9%). The ancestral “Manu” sublineages, Manu1 and Manu2, each comprised 11 isolates. Additionally, the T3-ETH sublineage of the EA lineage included nine isolates. Notably, no strains belonging to the Beijing or Haarlem sublineages were detected in this study.

**Table 1.** Summary of the spoligotyping results of the 116 samples with interpretable results.

| Lineage, sublineage (n) | SIT or Orphan (No. of isolates, % of the total isolates) |
| --- | --- |
| EA, T (n = 18) | SIT 53 (n = 13, 11.2%)<br>SIT 40 (n = 2, 1.7%)<br>SIT 334 (n = 1, 0.9%)<br>Orphan (n = 2, 1.7%) |
| EA, T3-ETH (n = 9) | SIT 149 (n = 9, 7.8%) |
| EA, T2 (n = 6) | SIT 52 (n = 3, 2.6%)<br>Orphan (n = 3, 2.6%) |
| EA, LAM5 (n = 1) | Orphan (n = 1, 0.9%) |
| EA, T3 (n = 6) | Orphan (n = 6, 5.1%) |
| EA, H3-Ural-1 (n = 1) | SIT 35 (n = 1, 0.9%) |
| EA, H3 (n = 2) | SIT 764 (n = 1, 0.9%)<br>SIT 50 (n = 1, 0.9%) |
| EA, LAM (n = 2) | SIT 42 (n = 2, 1.7) |
| EA, Turkey (n = 1) | Orphan (n =1, 0.9%) |
| EA, T1-RUS2 (n = 3) | SIT 4 (n = 3, 2.6%) |
| EAI, CAS1-Delhi (n = 16) | SIT 952 (n = 1, 0.9%)<br>SIT 26 (n = 6, 5.2%)<br>SIT 309 (n = 2, 1.7%)<br>SIT 25 (n = 3, 2.6%)<br>Orphan (n = 4, 3.4%) |
| EAI, CAS1-Kili (n = 2) | SIT 21 (n = 2, 1.7%) |
| EAI, CAS (n = 1) | Orphan (n = 1, 0.9%) |
| Lineage 7, ETH1 (n = 24) | SIT 910 (n = 21, 18.0%)<br>SIT 1729 (n = 3, 2.6%) |
| IO, Manu2 (n = 11) | SIT 54 (n = 9, 7.8%)<br>Orphan (n =2, 1.7%) |
| IO, Manu1 (n = 11) | SIT 523 (n = 7, 6.0%)<br>Orphan (n = 4, 3.4%) |
| IO, Manu ancestor (n = 1) | Orphan (n =1, 0.9%) |
| Unknown (n =1) | Orphan (n =1, 0.9%) |

EA: Euro-American; EAI: East-African-Indian; IO: Indo-Oceanic; SIT: Spoligotyping International Type

#### Shared and Orphan Strains

Thirty-nine distinct spoligotype patterns were identified among the 115 known Mtb strains. **Shared strains:** Among the 39 distinct spoligotypes, 19 (48.7%) were shared types comprising 90 strains recorded in the SITVIT2 database. The most common were SIT 910 (23.3%, ETH lineage), SIT 53 (14.4%, EA lineage), followed by SIT 54 (Manu 2) and SIT 149 (T3-ETH), each with 9 strains (10.0%) **(Fig 2)**

**Fig 2.**
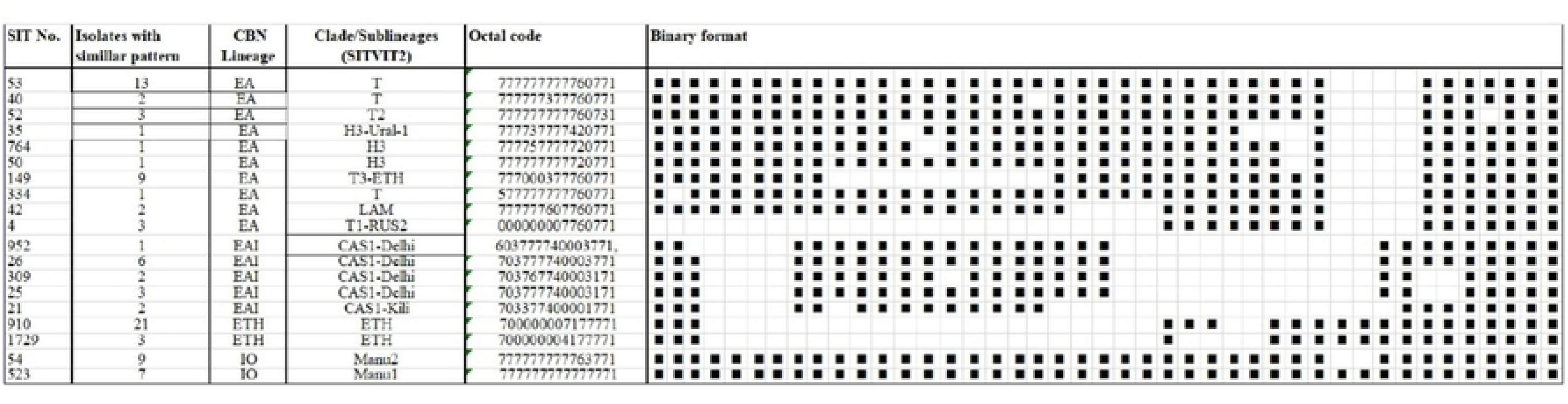
The spoligotype patterns of shared strains (n = 90) along with their corresponding lineages and sublineages identified from Mtb isolates from presumptive TB patients at two public hospitals in East Gojjam Zone, northwest Ethiopia. CAS: Central Asian; CBN: Conformal Bayesian Network; EA: Euro-American (L4); EAI: East African Indian (L3); ETH: Ethiopian (L7); IO: Indo-Oceanic (L1); L: Lineage; SIT: Spoligo-international type.

### Orphan Strains

25 strains, grouped into 20 unique patterns not found in the database, were classified as orphan strains (**Fig 3).**

**Fig. 3.**
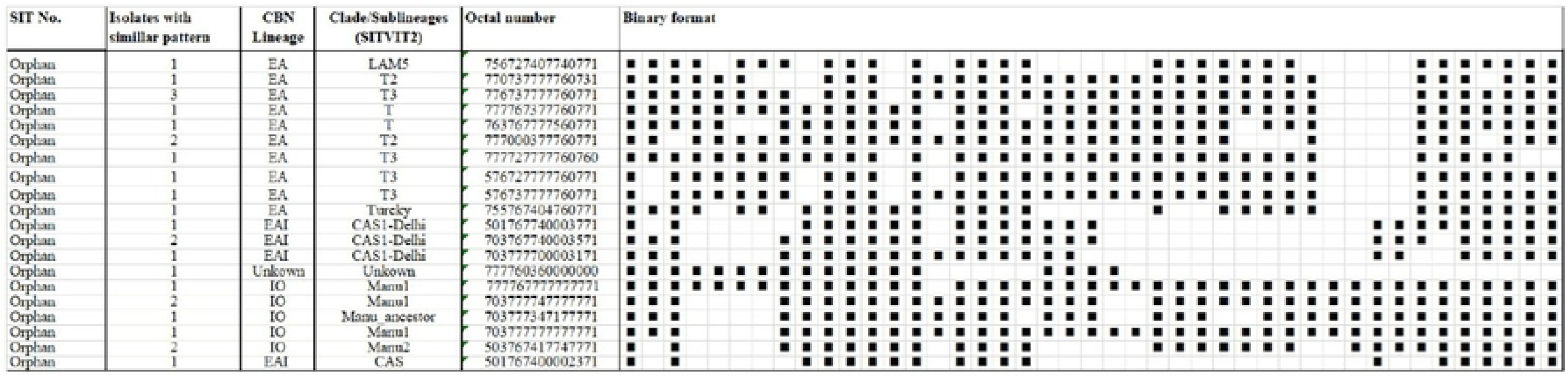
The spoligotype patterns of orphan strains (n = 25) along with their corresponding lineages and sublineages identified from Mtb isolates from presumptive TB patients at two public hospitals in East Gojjam Zone, northwest Ethiopia. CAS: Central Asian; CBN: Conformal Bayesian Network; EA: Euro-American (L4); EAI: East African Indian (L3); IO: Indo-Oceanic (L1); L: Lineage; SIT: Spoligo-international type; Orphan indicates strains not found in the database.

### Geographical distribution of sub-lineages

Geographic mapping was performed for 74 (64%) Mtb strains; the remaining lacked location data. The investigation revealed diverse TB strain distribution across woredas in the zone. Debre Elias had the most strains of any of the woredas, indicating a more diverse TB transmission environment. Strains such as CAS1-Kil, CAS1-Delhi, MANU2, and ETH1 were found here, indicating numerous sources or long-term transmission. In contrast, Aneded had a relatively low presence of TB strains, with just CAS1-Delhi being detected. Baso Liben has moderate strain variety, including CAS1-Delhi, MANU2, and H3-Ural-1, indicating that multiple lineages are being transmitted concurrently. Meanwhile, Enemay had fewer strains overall, particularly T3-ETH, ETH1, and LAM5, which could hint at limited transmission of a few dominant strains. Several strains, including ETH1 and H3-Ural-1, were found in multiple woredas, indicating either extensive transmission or shared infection origins (**Fig 4).**

**Fig. 4.**
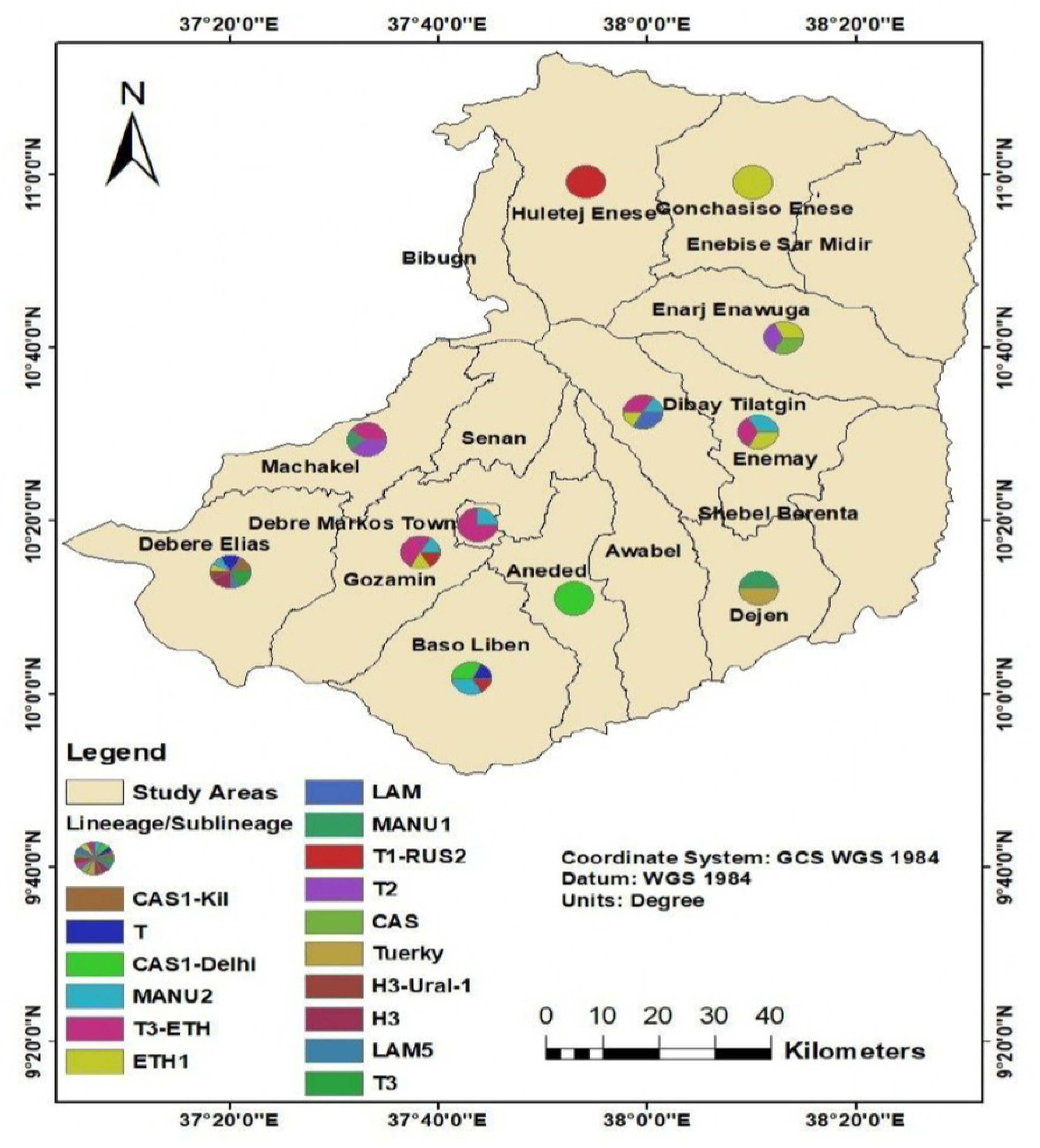
Geographic distribution of the study area and Mtb sublineages across Woredas in East Gojjam Zone (n = 67); pie charts show the proportion of each sublineage per Woreda. Source: Generated by the researcher using the East Gojjam Zone shapefile (2025).

### Clustering of identified strains

We have observed variation in the clustering rate across the major lineages and dominant sublineages, but overall, there was a high clustering rate in all cases. The majority, 96 (83.4%) of the isolates were clustered, whereas the remaining 20 (16.6%) were unique or singletons. Clustering was observed across the four major lineages identified. All 24 (100.0%) strains in the ETH lineage (lineage 7) and 20 (86.9%) of the strains in the IO lineage (lineage 1) were clustered. Moreover, 15 (78.9%) and 37 (75.5%) of the 49 strains were clustered within the EAI and EA lineages, respectively **(Table 2).**

**Table 2:** Clustering rate of Mtb strains recovered from sputum samples of patients attending two public hospitals in the East Gojjam Zone, Northwest Ethiopia (n = 115).

| Lineage | Number of isolates (n) | Clustered strains (n) | Clustering rate (%) | Cluster number (n) | Cluster size (n) | SIT | Sublineage |
| --- | --- | --- | --- | --- | --- | --- | --- |
| EA (L4) | 49 | 37 | 75.5% | 8 | 13 | 53 | T |
|  |  |  |  |  | 2 | 40 | T |
|  |  |  |  |  | 3 | 52 | T2 |
|  |  |  |  |  | 3 | Orphan | T3 |
|  |  |  |  |  | 2 | Orphan | T2 |
|  |  |  |  |  | 9 | 149 | T3-ETH |
|  |  |  |  |  | 2 | 42 | LAM |
|  |  |  |  |  | 3 | 4 | T1-RUS2 |
| ETH (L7) | 24 | 24 | 100.0% | 2 | 21 | 910 | AFRI |
|  |  |  |  |  | 3 | 1729 | AFRI |
| EAI (L3) | 19 | 15 | 78.9% | 5 | 6 | 26 | CAS1-Delhi |
|  |  |  |  |  | 2 | Orphan | CAS1-Delhi |
|  |  |  |  |  | 2 | 309 | CAS1-Delhi |
|  |  |  |  |  | 3 | 25 | CAS1-Delhi |
|  |  |  |  |  | 2 | 21 | CAS1-Kili |
| IO (L1) | 23 | 20 | 86.9% | 4 | 9 | 54 | Manu2 |
|  |  |  |  |  | 7 | 523 | Manu1 |
|  |  |  |  |  | 2 | Orphan | Manu1 |
|  |  |  |  |  | 2 | Orphan | Manu2 |

EA: Euro-American; EAI: East-African-Indian; ETH: Ethiopian; LAM: Latin-American-Mediterranean; L: Lineage; SIT: Spoligotype international type.

### Factors associated with clustering

Clustering rates were 100% for several sublineages, including ETH1 of lineage 7 (24 strains), CAS1-Delhi of lineage 3 (3 strains), T3-ETH of lineage 4 (9 strains), and Manu 2 of lineage 1 (11 strains). High clustering was also observed in the T and T2 sublineages, with 83.3% of strains clustering in each. No statistically significant association was found between clustering and drug resistance (p = 0.61) **(Table 3).**

**Table 3:** Clustering by major Mtb lineages, sublineages, and HIV status (n = 115).

| Characteristics | Total<br>n (%) | Clustered<br>n (%) | Non-clustered<br>n (%) | P value |
| --- | --- | --- | --- | --- |
| **Major lineages** |  |  |  | 0.03* |
| EA (L4) | 49 (42.6) | 37 (75.5) | 12 (24.5) |  |
| EAI (L3) | 19 (16.5) | 15 (78.9) | 4 (21.1) |  |
| IO (L1) | 23 (20.0) | 20 (86.9) | 3 (13.1) |  |
| ETH (L7) | 24 (20.9) | 24 (100.0) | 0 (0.0) |  |
| **Sublineage** |  |  |  | <0.001* |
| ETH1 | 24 (20.9) | 24 (100.0) | 0 (0.0) |  |
| CAS1-Delhi | 16 (13.9) | 3 (100.0) | 0 (0.0) |  |
| Manu1 | 11 (9.6) | 9 (81.8) | 2 (18.2) |  |
| Manu2 | 11 (9.6) | 11 (100.0) | 0 (0.0) |  |
| T | 18 (15.7) | 15 (83.3) | 3 (16.7) |  |
| T2 | 6 (5.2) | 5 (83.3) | 1 (16.7) |  |
| T3 | 6 (5.2) | 3 (50.0) | 3 (50.0) |  |
| T3-ETH | 9 (7.8) | 9 (100.0) | 0 (0.0) |  |
| Others<sup>#</sup> | 14 (12.1) | 7 (50.0) | 7 (50.0) |  |
| **Any resistance** |  |  |  | 0.61 |
| Yes | 18 (20.2) | 16 (88.9) | 2 (11.1) |  |
| No | 71 (79.8) | 62 (87.3) | 9 (12.7) |  |

\*: Fisher's exact test; #: CAS (n = 1); CAS1-Kili (n = 2); H3 (n = 2); H3-Ural-1 (n = 1); LAM (n = 2); LAM5 (n = 1); L: Lineage; Manu ancestor (n = 1); T1-RUS2 (n = 3); Turkey (n = 1)

## Discussion

In this study, four major Mtb lineages were identified, with the EA lineage being the most prevalent (42.6%), followed by Lineage 7 (20.8%), the OI lineage (20.0%), and the EAI lineage (16.5%). A total of 17 sublineages were identified within these lineages. The ETH sublineage of Lineage 7 was the most common (20.8%), followed by the ill-defined T clade of EA (15.6%) and the CAS1-Delhi sublineage of EAI (13.9%). No strains belonging to the Beijing or Haarlem clades were found. Among the 115 isolates analyzed, 39 distinct spoligotype patterns were observed. Of these, 19 patterns (48.7%) matched known shared international types (SITs) in the SITVIT2 database, while 20 (51.3%) patterns were classified as orphan strains. The most frequently observed SITs were SIT 910 (23.3%) from Lineage 7 and SIT 53 (14.4%) from EA. A considerable proportion of SIT 54 (10.0%) from the Manu 2 sublineage was also detected. Overall, 96 isolates (82.7%) were grouped into clusters, indicating potential recent transmission, while 20 isolates (17.3%) were unique.

Ethiopia harbors a genetically diverse population of Mtb, encompassing both ancient and modern lineages. The four lineages identified in this study have also been consistently reported in previous research across various regions of the country, with the EA lineage frequently emerging as the most prevalent (9). In our study, the EA, which consisted of nine different sublineages, accounted for 42.6% of strains, which is lower than proportions reported in the Arsi Zone of southeastern Ethiopia (79.3%) (18), a multi-zone study in the Oromia region (79.4%) (19), data from other parts of northwestern Ethiopia (55.2%) (20), and pooled findings from a systematic review (64.8%) (9). However, our figure (42.6%) is slightly higher than that reported in the Gondar area of northwestern Ethiopia (40.1%) (21). Although not as highly related to hypervirulence as Lineage 2 (Beijing) strains, EA strains contribute significantly to the worldwide TB burden. Their widespread geographic distribution is most likely due to their relatively high transmissibility and the mobility of individuals between surrounding regions, which facilitates their continuous spread. The dominance of this lineage has substantial public health implications, particularly since several of its sublineages have been related to outbreaks of MDR TB (22, 23).

The proportions of Lineage 7 (20.8%) and IO lineage (20.0%) observed in our study were higher than those reported previously. Lineage 7 was initially discovered in the Weldiya area of northeastern Ethiopia, where 17 of the 133 isolates (12.7%) were found to be Lineage 7 (24). Subsequent studies from different parts of northwest Ethiopia documented Lineage 7 prevalence ranging from 1.5 to 15.6% (20, 24–26), and a systematic review noted that Lineage 7 accounts for 4.1% of Mtb strains in Ethiopia (9). The higher proportion observed in our study may be attributed to the geographic location of our study site, which experiences population movement from both northeastern and northwestern Ethiopia, potentially facilitating increased transmission of this lineage. Notably, Lineage 7 has been associated with delayed healthcare-seeking behavior among TB patients, which could contribute to ongoing community transmission and pose important public health concerns (27). Further investigation using advanced molecular tools such as whole-genome sequencing is warranted to confirm these findings.

The proportion of the IO lineage, represented by the Manu sublineages, observed in this study (20.0%) is notably higher than previously reported in various regions of Ethiopia. For instance, a study from the Oromia region reported a prevalence of 9.8%, while studies from the Amhara region documented lower rates ranging from 2.6% to 5% (20, 24, 25), with a pooled prevalence of 7.1% identified in a systematic review (9). The IO lineage is recognized as the predominant strain causing TB in South Asian populations (28). However, a higher prevalence of the IO lineage (27.15%) was reported in Egypt in North Africa (29). Its presence in the current study area suggests a possible ancient introduction followed by localized transmission. Although strains within the IO lineage are generally considered less transmissible and less virulent compared to modern lineages such as Beijing and EA, they have been associated with latent infections and slower disease progression (30). These characteristics can pose challenges for public health strategies that prioritize active case detection. Therefore, effective TB control in regions where this lineage is prevalent may require targeted approaches, including tailored diagnostics, robust contact tracing, and proactive management of latent TB infections.

The EAI lineage is among the most frequently identified Mtb lineages in Ethiopia. A study conducted in the Gondar area of northwest Ethiopia reported it as the dominant strain, accounting for 53.6% of cases, surpassing even the EA lineage (21). Other studies have identified the EAI lineage as the second most prevalent, with reported prevalences ranging between 9.8% and 42.2% (18, 20, 31). A systematic review further estimated its overall prevalence at 23.0% (9). In the current study, 16.5% of the isolates belonged to the EAI lineage, aligning with the general range reported in earlier studies, though it emerged as the least common strain in this specific area. While the EAI lineage tends to exhibit reduced potential for rapid transmission compared to more modern lineages such as Beijing and EA, recent reports of emerging drug resistance underscore the need for continuous monitoring and surveillance (32, 33).

The predominant sub-lineage identified in this study was ETH1 of Lineage 7, with the most common shared type being SIT 910 (18.0%) within the same lineage. This may suggest the successful local expansion of this particular clone, potentially indicating a highly fit strain circulating in the study area. According to a systematic review, the most common Mtb sublineages in Ethiopia are the T family (48.0%), CAS (23.0%), Haarlem (11.0%), Manu (6.0%), and ETH (4.1%) (9). We reported most of these sublineages in the current study area, with the ill-defined T superfamily (15.6%) and CAS1-Delhi sublineage (13.6%) being the second and third most prevalent, respectively. However, it is important to note that the T family is not a monophyletic group; it does not represent a single evolutionary lineage. Rather, it serves as a default category for strains that cannot be definitively assigned to well-defined lineages such as Haarlem, LAM, CAS, or EAI (34). The CAS1 Delhi of the EAI lineage was also reported as the predominant sub-lineage from previous studies in the Gondar area of northwest Ethiopia (21). One of the most clinically and epidemiologically significant Mtb lineages in the world is the Beijing sublineage, which is a member of Lineage 2 (East Asian lineage). It is widely distributed and is particularly known for its strong association with drug resistance, elevated transmissibility, and ability to evade host immune responses—all of which pose major challenges to TB control and effective patient management (35). However, in our study, this sublineage was not detected. Consistent with our findings, the Beijing lineage has been infrequently reported in previous studies conducted in Ethiopia (23, 36), with a systematic review indicating it accounts for only 0.2% of Mtb strains in the country (9).

A systematic review documented SIT149 as the most common shared type, followed by SIT53 and SIT25 (9). In our study, SIT 53 (14.4%) of the EA lineage was the second most prevalent shared strain, whereas SIT 54 (Manu 2 sublineage) and SIT 149 (T3-ETH sublineage) were equally the third most prevalent shared types. A rare spoligotype pattern of the SIT41 of Turkey sublineage has been detected from one of the districts in the study area, and previously, this strain has been reported after the report from the North Shewa Zone of the Amhara region and in Gondar (20, 25). The proportion of shared Mtb strains in Ethiopia has been reported to range from 0.5% in the Harari region to 36% in the Amhara region (9). In this study, 48.7% of the isolates were identified as shared types, higher than previously reported figures. However, 51.3% of the strains remain orphan types, underscoring the need for further investigation using more advanced molecular techniques. Such studies could discover additional, possibly novel, strains circulating in the area.

The overall clustering rate of 82.7% observed in our study is notably higher than previously reported rates, including 52.4% in the Oromia region (19), 70% in the Ethiopian national survey (31), and 76.2% in the Afar region (37). This high clustering rate indicates an extensive level of recent transmission of specific Mtb clones in the study area, particularly the strains belonging to ETH 1, CAS1-Delhi, and Manu 2 sublineages, which exhibited clustering rates of 100%. Conducting further research using advanced molecular techniques combined with Geographic Information System (GIS) tools is essential to gain a more comprehensive understanding of the Mtb transmission dynamics in the area.

This study offers valuable insights into the genetic diversity of Mtb strains in a rural area of Ethiopia, although it has certain limitations. One major constraint is the reliance on spoligotyping, which, despite its utility, offers limited resolution for distinguishing some lineages. While enhanced methods such as MIRU-VNTR or whole genome sequencing offer greater discriminatory power, they are often inaccessible in low-resource settings due to financial and infrastructural limitations (10–12). Another concern is the over-decade gap between the initial isolation of strains and their molecular analysis, which may raise questions about the current relevance of the data. The delay was primarily attributed to financial and logistical challenges. Given the low mutation rate of Mtb (∼0.3–0.5 mutations per genome per year), the genetic stability of the strains supports the continued relevance of the findings (38). As the first molecular epidemiological study from this region, these results provide an essential reference point for future research and TB control strategies.

## Conclusion

This study highlights considerable genetic diversity among Mtb strains in a rural Ethiopian setting, with four major lineages identified—EA, Lineage 7, IO, and EAI. The unexpectedly high proportions of Lineage 7 and IO strains, coupled with a high clustering rate (82.7%), indicate active local transmission, particularly of the ETH1, CAS1-Delhi, and Manu2 sublineages, and underscore the importance of implementing tailored TB control strategies in the area. The presence of many orphan spoligotypes points to unique or underreported strains, emphasizing the need for advanced molecular tools like whole genome sequencing.

## Acknowledgments

The authors gratefully acknowledge the AHRI for its generous support, both financially and materially. We also extend our thanks to the laboratory professionals at Debre Markos Referral Hospital and Mota District Hospital, with special appreciation to Mr. Derejew Zewdie for his significant contributions to data collection.

## Notes

### Competing Interest Statement

The authors have declared no competing interest.

